# Campolina: A Deep Neural Framework for Accurate Segmentation of Nanopore Signals

**DOI:** 10.1101/2025.07.08.663658

**Authors:** Sara Bakić, Krešimir Friganović, Bryan Hooi, Mile Šikić

## Abstract

Nanopore sequencing enables real-time, long-read analysis by processing raw signals as they are produced. A key step, segmentation of signals into events, is typically handled algorithmically, struggling in noisy regions. We present Campolina, a first deep-learning framework for accurate segmentation of raw nanopore signals. Campolina uses a convolutional model to identify event boundaries and significantly outperforms the traditional Scrappie algorithm on R9.4.1 and R10.4.1 datasets. We introduce a comprehensive evaluation pipeline and show that Campolina aligns better with reference-guided ground-truth segmentation. We show that integrating Campolina segmentation into real-time frameworks, Sigmoni and RawHash2, improves their performance while maintaining time efficiency.

## 1 Background

Nanopore sequencing [1] has emerged as a transformative technology in genomics, offering realtime, long-read sequencing capabilities that enable direct analysis of nucleic acids. As nucleotides pass through a nanopore, they generate disruptions in the ionic current. These disruptions are measured, producing raw nanopore signals, and processed to extract meaningful biological information. The most common practice is to translate raw nanopore signals into a sequence of nucleotides through basecalling, the decoding of the nucleotide sequence, and process the basecalled sequence for various tasks [2]. Oxford Nanopore Technology (ONT) has developed several deep learning-based basecallers [3, 4]. Their growing complexity creates a computational time bottleneck, which motivates a shift towards the development of methods for direct processing of raw nanopore signals in real-time. Computational methods for analyzing raw nanopore signals at the same rate at which they are produced during nanopore sequencing have several advantages, including parallel execution of sequencing and analysis, and the early termination of a read or an entire sequencing run without sequencing the full molecule or sample, through techniques known as adaptive sampling [5], Read Until [6] and Run Until [7]. These techniques can trigger a signal for ejecting the current read from the nanopore once a certain condition is satisfied, i.e., a target species abundance in a metagenomic sample is reached, or a region of interest is not matched [8–10]. Developing accurate and efficient real-time analysis techniques enables reductions in both time and cost associated with genome analysis.

RawHash2 [11, 12], Sigmoni [13], and UNCALLED [14] are methods developed to process raw nanopore signals for different downstream tasks without the need for basecalling. RawHash2 is a hash-based mechanism designed to map raw nanopore signals to reference genomes in real time. It generates an index from the reference genome and efficiently maps raw signals by matching their hash values. This method enables rapid and accurate alignment, even for large genomes, by employing adaptive quantization and chaining algorithms. Sigmoni is a tool focused on the classification of nanopore signals using a compressed pangenomic index. It leverages advanced indexing techniques to classify raw nanopore signals efficiently, facilitating applications such as pathogen detection and metagenomic analysis. UNCALLED is a read mapper that aligns raw nanopore signals directly to DNA references, enabling real-time targeted sequencing. It allows for software-based adaptive sampling on platforms like the Oxford Nanopore MinION ^1^ or GridION ^2^, providing flexibility in sequencing workflows.

The advancements proposed in the aforementioned frameworks reflect a growing trend toward analyzing raw signals in real-time, aiming to reduce computational overhead and improve the efficiency of genomic analyses.

A critical step in the raw signal-processing algorithms is the segmentation of raw signals into discrete “events”, which correspond to the translocations of individual nucleotides through the nanopore. Frameworks such as Scrappie [15], developed by ONT, have been widely used for this purpose. Scrappie detects events by performing a rolling t-test to identify significant changes in picoamp (pA) levels and groups stretches of signal into events. However, this approach relies heavily on a predefined set of hyperparameters, such as window length, peak height, and minimum inter-peak distance.

The statistical event detection setup combined with the predefined, fixed set of hyperparameters and thresholds is not robust to the diverse characteristics of nanopore signals, which can exhibit significant variations in signal-to-noise ratios, event lengths, and current level distributions. Consequently, Scrappie often suffers from over-segmenting or failing to retrieve events in noisy regions, leading to errors that propagate through the downstream analysis pipeline. The nontriviality of finding optimal and robust hyperparameters for Scrappie segmentation is mirrored in the fact that, despite relying on the same segmentation algorithm, different methods for raw signal processing define different sets of segmentation hyperparameters. Furthermore, the existing pipelines typically introduce additional postprocessing of the obtained segmentation, such as compressing same-character runs, to correct for segmentation errors caused by oversegmenting signals in noisy regions. The constant development of new nanopores that produce signals with different characteristics aggravates this problem even further. From the abovementioned methods, only RawHash2 provides a set of segmentation-specific hyperparameters specifically designed for the most recent R10.4.1 pore version. Other frameworks only provide an R9.4.1 setup, which is considered deprecated.

The Remora framework [16], developed as part of the Dorado basecalling ecosystem, has recently introduced a signal refinement step to adjust signal scaling and mappings to improve matching to the expected k-mer levels. Remora combines signal-to-sequence alignment with precise signal segmentation, providing a high-quality ground truth for evaluating segmentation models. This is achieved with several steps: basecalling signals with Dorado, aligning the basecalled sequence to the provided reference with minimap2 [17], and refining the segmentation using signal-to-sequence mapping anchored on the reference.

While there are established approaches to generating accurate signal segmentation based on refining the signal scaling and the signal-to-sequence mapping to more closely match the expected signal levels [18], these approaches require basecalling and mapping steps as part of the pipeline, and are, as such, not suited for real-time processing. On the contrary, the only approach to direct segmentation of the nanopore signals is Scrappie, which relies on sets of predefined hyperparameters and thresholds and is not robust in noisy regions.

To address these challenges, we propose Campolina, a novel framework for nanopore signal segmentation. Campolina is a deep learning-based architecture that outputs accurate segmentation of raw nanopore signals without basecalling, alignment, or refinement. Campolina does not rely on a predefined set of hyperparameters or reference for alignment, and is more robust than the traditional algorithmic approaches. We assess the quality of the obtained segmentation and demonstrate its suitability for existing raw signal processing frameworks. We devised an evaluation pipeline that truthfully measures the quality of nanopore signal segmentation and analyzed the benefits of Campolina segmentation in existing pipelines that rely on traditional algorithmic approaches. We performed tests on various datasets sequenced with R9.4.1 and R10.4.1 pore versions and in various downstream setups, showing that Campolina segmentation improves segmentation quality and is well-suited for existing frameworks that process raw nanopore signals. To summarize, our contributions in this paper are as follows:

- We introduce Campolina, a lightweight deep neural framework that processes raw nanopore signals to produce high-quality segmentation. Campolina improves the quality of the obtained segmentation when compared to the traditional algorithmic approaches and mitigates the dependency on predefined sets of thresholds and references. We provide Campolina models for the R9.4.1 and R10.4.1 nanopore versions.
- We propose a segmentation evaluation pipeline that assesses segmentation quality beyond simple set matching with respect to ground-truth segmentation and the expected values from the corresponding reference sequence. The proposed evaluation ensures a complete and truthful assessment of the segmentation quality by looking into the predicted border positions and the obtained event levels.
- We analyze the suitability of Campolina segmentation in the existing frameworks that analyze raw nanopore signals for various classification tasks. We show that Campolina’s segmentation improves the classification accuracy across different frameworks, nanopore versions, and datasets when compared to the traditional segmentation approach.

## 2 Results

### Campolina framework for accurate segmentation of nanopore signals

Campolina processes a time-series of fixed length *L* augmented point-wise with four statistical descriptors (Figure 1(a)) with the objective of predicting positions in the input that correspond to an event border. Campolina (Figure 1(b)) consists of a convolutional subnetwork [19] that processes the input signal to extract semantic features, and a classification head that outputs a non-normalized probability (logit) for each input point indicating whether the point is a border or not. The model is trained in a supervised manner with ground truth border positions extracted based on a two-step ground truth pipeline shown in Figure 2. In the first step, the signal is basecalled with a “super accurate” Dorado basecaller and aligned to the reference with minimap2, which is incorporated in the Dorado framework. Along with the basecall and mapping information, the initial approximation of signal segmentation in the form of a move table is extracted. The output of the first step, along with the k-mer model, is passed to the Remora refinement step, from which the accurate event border positions are obtained.

**Figure 1.**
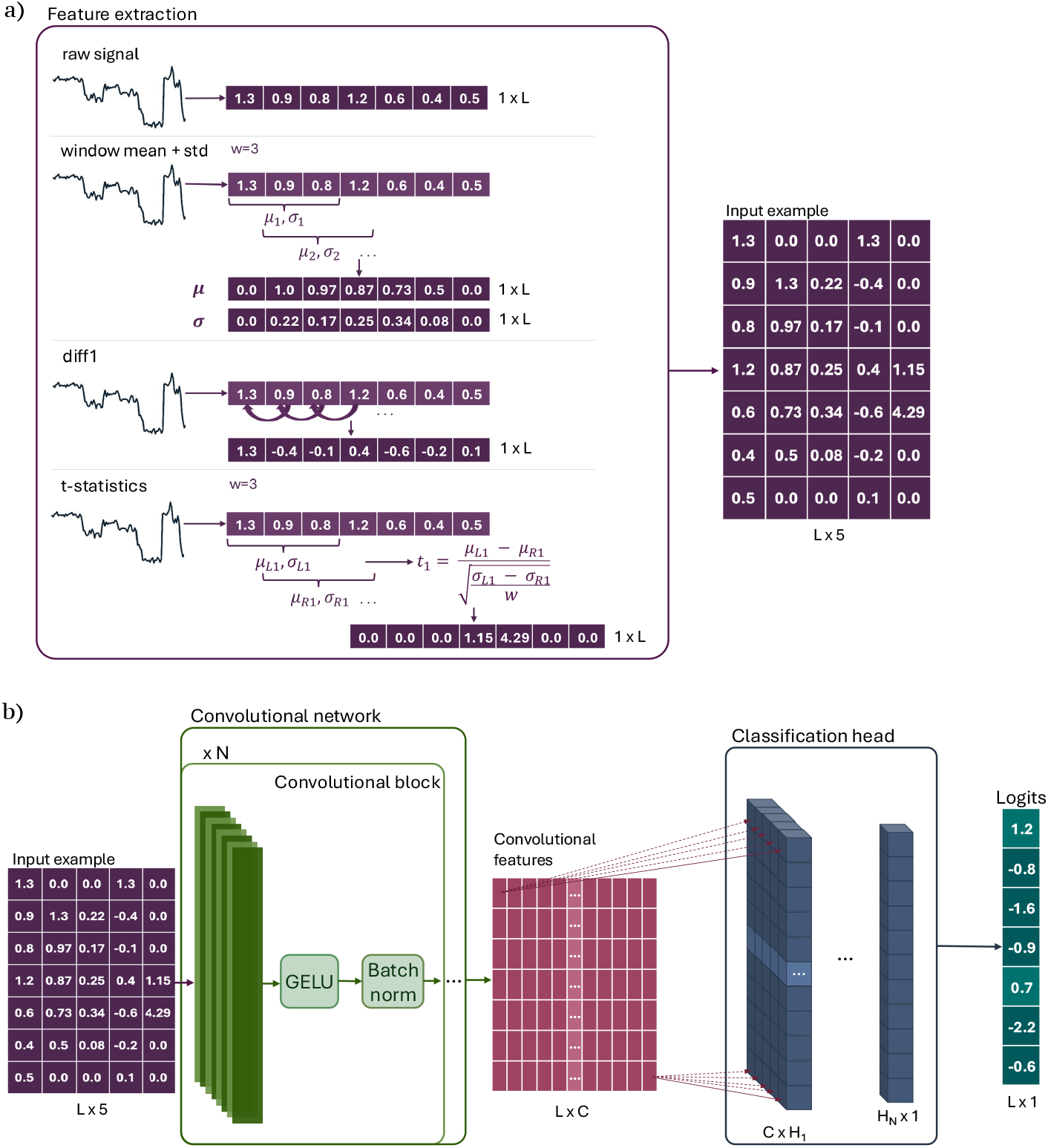
Full Campolina framework. a) **Feature extraction pipeline:** Raw signals are divided into non-overlapping chunks of fixed length *L* = 6000 samples. Each chunk is z-normalized and augmented with four descriptors, rolling window mean and standard deviation (*std*), first-order difference (*diff1*), and rolling-window *t-statistics*, resulting in a 5-channel input representation per chunk. b) **Model architecture:** The processed input is passed through a convolutional network consisting of five convolutional blocks with increasing channel dimensions (32, 64, 64, 128, 128), kernel size 3 in the first block and 31 in subsequent blocks, and GELU activations. A linear classification head outputs a non-normalized probability for each point, predicting the likelihood of an event border. The final output has shape 6000 *×* 1.

**Figure 2.**
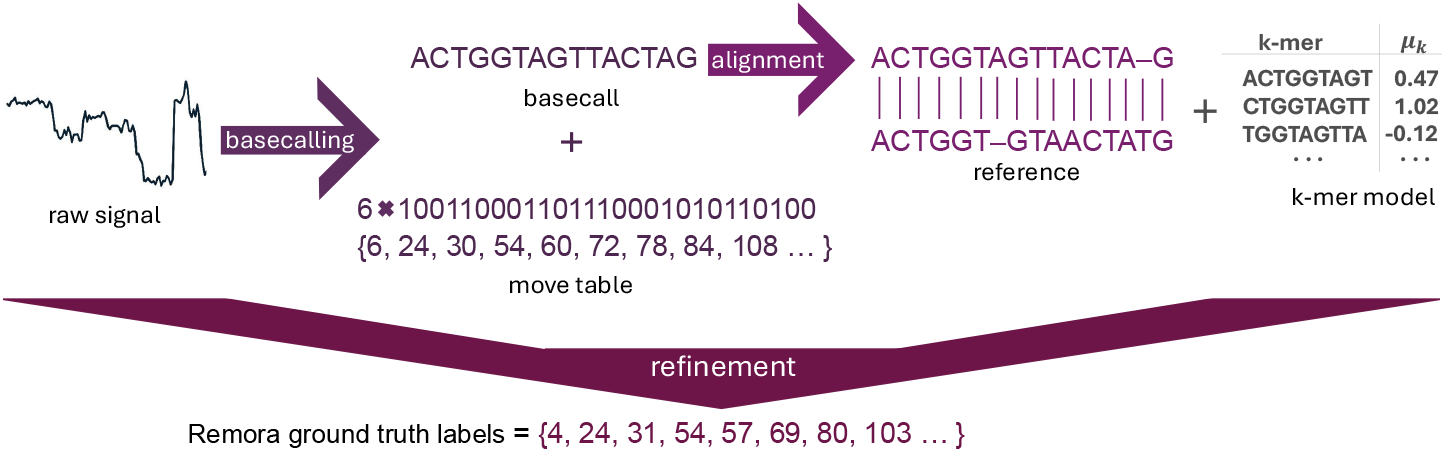
Ground truth pipeline. In the first step, raw nanopore signals are basecalled with the Dorado basecaller, which internally uses minimap2 to align the reads to a reference genome. This step also produces a move table that provides an initial approximation of signal segmentation. The resulting basecall, alignment details, and move table, together with a k-mer model that describes expected k-mer levels, are then input to the Remora refinement step to obtain accurate event boundary positions.

Campolina is trained by optimizing a three-component composite loss. The accuracy of identifying correct event borders is optimized through Focal loss to account for the prevalence of negative labels, i.e., positions that do not correspond to an event border. Huber loss component constrains the number of predicted event borders given the number of borders in the labels. Finally, we define a custom Consecutive loss component that prevents the model from predicting consecutive points in the signal as event borders, which is especially significant for the consecutive points measured as the nucleotides in the nanopore are shifting. The contribution of each loss component is quantified in the ablation study, where we trained additional models after removing certain loss components. The details of the ablation study are available in Methods and Supplementary Material S2.

We train and evaluate the R9.4.1 and R10.4.1 versions of Campolina, which is, to the best of our knowledge, the first deep learning based framework for direct segmentation of raw nanopore signals. Moreover, since the traditional algorithmic segmentation was initially developed for the R9.4.1 nanopore version, Campolina is the first tool developed specifically for segmenting signals sequenced with the contemporary R10.4.1 nanopore version.

### Need for comprehensive assessment of segmentation quality

A naive approach to evaluating segmentation by simply comparing the sets of predicted and ground truth borders is insufficient for a truthful assessment of segmentation quality. Multiple measurements can be taken during the event change, which is why the effects of local perturbations to the border positions on real-time signal analysis are unclear. We inspect the effects of perturbations to the event border positions by applying local shifts, deletions, or insertions of the ground truth borders and inspecting the effects on multiclass classification accuracy (Figure 3(a)). We use the border positions obtained through the refinement offered in the Remora framework and incorporate the perturbed borders into the raw signal processing framework RawHash2.

**Figure 3.**
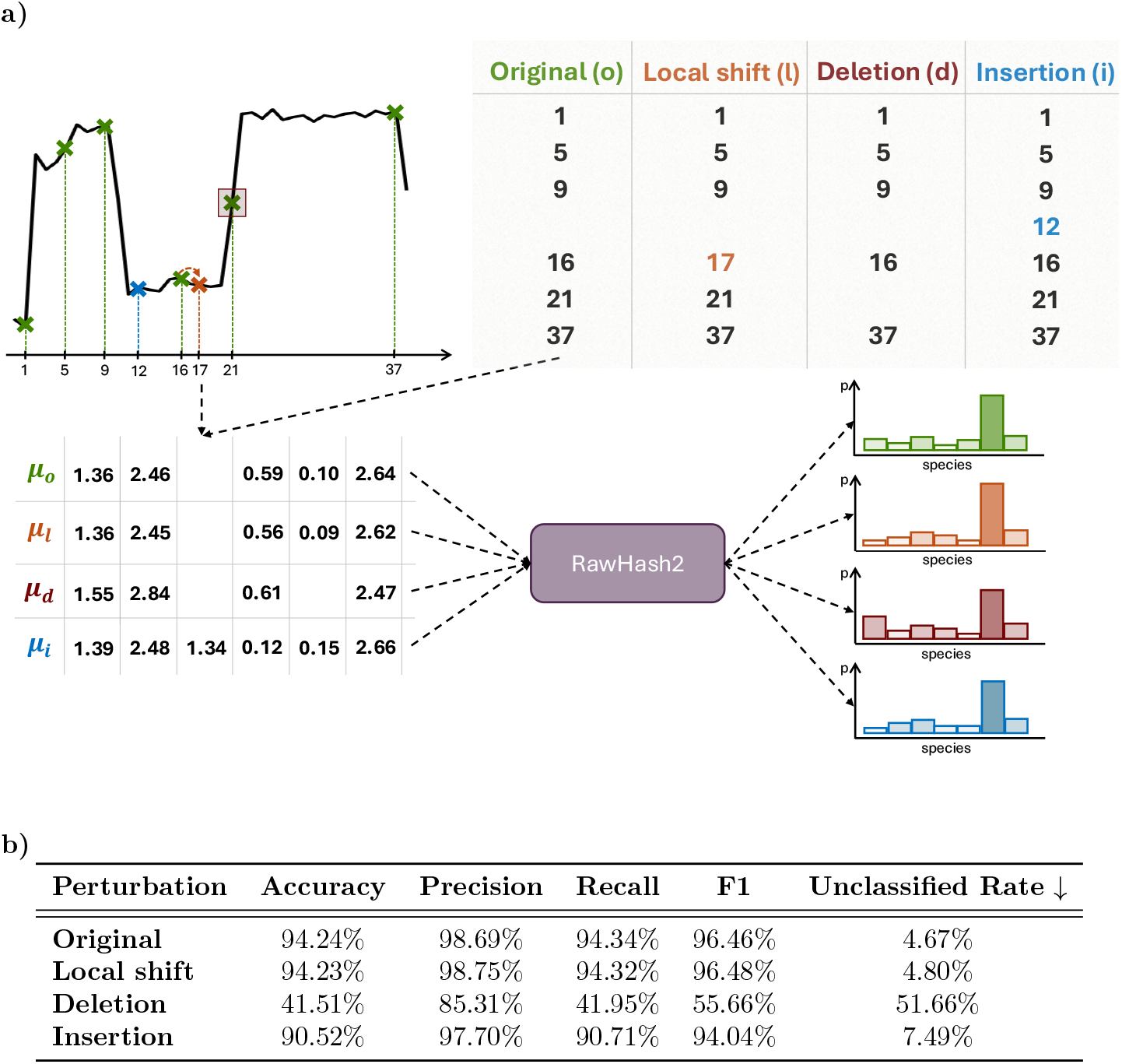
The effects of border perturbations. The performance of the RawHash2 framework on a Zymo multi-class classification task is inspected for the refined Remora borders, and three perturbed versions where a fraction of borders is shifted (local shift), deleted (deletion), or a fraction of events is split into two (insertion) as illustrated in **a).** We report the accuracy, precision, recall, F1 score, and unclassified rate in **b)**. Unclassified rate is the percentage of reads that could not be assigned to any of the provided references due to poor hash matching. We treat those reads as misclassified when calculating other metrics, but also provide the details on the “unclassified rate” to quantify the ratio of reads that cannot be aligned to any reference.

#### Local shift

In the first experiment, we randomly choose 20% of the ground truth borders and shift the border position randomly to one position left or right. This perturbation does not affect the number of segments in the signal but imposes small changes to the normalized means of the obtained events.

#### Deletion

In the second experiment, we delete 20% of randomly chosen borders. This perturbation reduces the number of predicted events, and by merging two or more events into a single event, it affects the normalized means of the obtained events.

#### Insertion

In the third experiment, we randomly choose 20% of the ground truth events that we split into two events of the same length by inserting a border in the middle of the selected event. This perturbation increases the number of predicted events and slightly affects the normalized means of the events. The effects to the normalized event levels are not as strong as in the deletion case, since the inserted events have normalized means similar to the levels of the original event.

Figure 3(b) shows the results obtained in the R10.4.1 Zymo multi-class classification task after performing local shifts, deletions, and insertions of the event borders. The results show that local shifts of the border positions have no negative effect on the downstream performance. On the other hand, deleting and inserting borders downgrades the classification performance. While the deletion effect is very strong, the insertion effect is less obvious, arguably due to the RawHash2 relying on a segmentation post-processing tailored for the Scrappie algorithm, which is prone to over-segmenting in noisy regions and, therefore, resolving some issues related to over-segmentation introduced by adding event borders. Nonetheless, the perturbation effect on results shows that the evaluation of the segmentation quality needs to surpass a simple border matching approach to ensure a truthful assessment of the segmentation. In this paper, we propose an evaluation pipeline that, beyond comparing the border positions, quantifies how well the predicted segmentation corresponds to the ground truth, and how well the obtained event levels match the ground truth and the expected levels based on the reference.

### Campolina improves the quality of segmentation over the traditional algorithmic approach

To assess the quality and suitability of Campolina segmentation, we conducted a systematic analysis using two datasets sequenced with R9.4.1 and R10.4.1 pore versions. The first dataset is ZymoBIOMICS HMW DNA Standard (D6322), which consists of a mixture of high molecular weight genomic DNA isolated from pure cultures of seven bacterial and one fungal strains. The species included in this analysis are: *S. aureus, S. enterica, P. aeruginosa, L. monocytogenes, E. faecalis, B. subtilis*, and *S. cerevisiae*. As *E. Coli* was used to train the Campolina border prediction model, we excluded those reads from the evaluation for possible bias. The second dataset is the B-lymphocyte cell line NA12878.

To evaluate the quality of segmentation, we systematically compare the segmentation obtained with Campolina with the ground truth segmentation as shown in Figure 4(a). The predicted border positions are initially compared with the ground truth ones, and Jaccard similarity between the two sets is calculated. Additionally, the predicted border positions are compared with the immediate neighbourhood of the ground truth positions ±1, and Jaccard similarity between these two sets is calculated and labeled as expanded Jaccard similarity. To further assess how well the predicted borders match the ground truth ones, we calculate the bi-directional Chamfer distance between the sets of predicted and ground truth positions. However, since we deem these metrics not sufficient for a complete assessment of the segmentation quality, we develop an evaluation pipeline in which we align the predicted segmentation to the ground truth one based on stretches of signal covered by a specific event pair.

**Figure 4.**
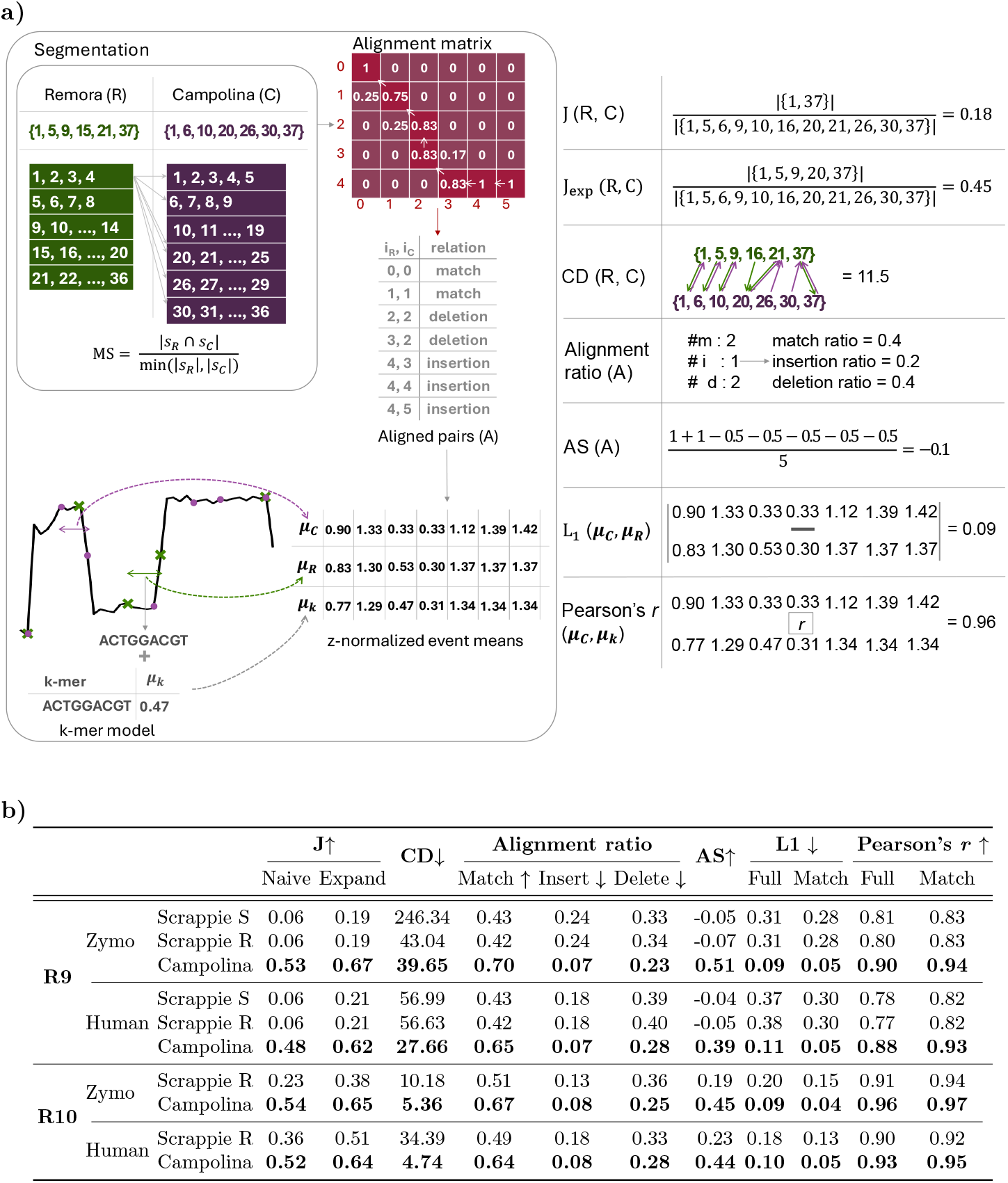
Segmentation quality evaluation. **a)** The predicted (C) and ground truth border positions (R) are compared using Jaccard similarity (J) and an expanded version that considers *±*1 positions (J_exp_). To further assess border alignment, the bi-directional Chamfer distance (CD) between positions (R and C) is computed. The event-based alignment by comparing predicted and ground truth events based on index overlap (MS), constructing an alignment matrix and performing traceback to identify alignment types: match (m), insertion (i), and deletion (d). Alignment ratios and alignment score (AS) are computed based on aligned pairs (A). The L1 distance between z-normalized means of aligned events (*µ*_*C*_ and *µ*_*R*_) is used to quantify event-level accuracy. Finally, reference k-mers are assigned to predicted events using the refined ground truth, and Pearson’s r is computed between predicted event means (*µ*_*C*_) and expected k-mer levels (*µ*_*k*_). **b)** The segmentation quality evaluation for R9.4.1 Zymo D6322 dataset, R9.4.1 Human NA12878 dataset, R10.4.1 Zymo D6322 dataset, and R10.4.1 Human NA12878 dataset. Both Zymo datasets are prepared without *E. coli* reads that were used for training. Segmentation results are compared against the segmentation obtained by the Scrappie algorithm with the hyperparameter set defined by Sigmoni (denoted as Scrappie S) and RawHash2 (denoted as Scrappie R). All metrics are calculated as shown in a). L1 and Pearson’s r are calculated for all alignments and matches only.

The process involves segmenting the signal into events, then comparing predicted and groundtruth events based on their index overlap to determine how well they match. An alignment matrix is constructed using these comparisons, with each element in the matrix representing a ratio of indices covered by ground truth and predicted event pair. We perform a traceback through the matrix and based on the traceback path, define three alignment types: *match* (m) when one predicted event aligns to one ground truth event, *insertion* (i) when multiple predicted events align to one ground truth event, and *deletion* (d) when one predicted event aligns to multiple ground truth events. Based on the alignment, we find ratios of ground truth events in match, insertion, and deletion alignment and calculate the alignment score. Furthermore, we calculate the L1 distance between z-normalized means of the aligned events for all alignments and only matches to quantify how well the obtained event levels match the ground truth ones. Finally, based on the event alignment and the alignment of the refined ground truth to the reference, we extract the corresponding reference k-mer for each predicted event, and based on the kmer model, calculate Pearson’s correlation coefficient *r* between the z-normalized mean of the predicted event and the expected k-mer level. Pearson’s correlation coefficients *r* are calculated for every alignment (Full), and for only matches (Match). A detailed explanation of the proposed evaluation pipeline is available in Methods.

We extract segmentation for all datasets with Campolina and Scrappie. For the Scrappie segmentation, we utilize sets of hyperparameters proposed in Sigmoni and RawHash2. While both RawHash2 and Sigmoni provide custom sets of segmentation hyperparameters for the R9.4.1 pore version, only RawHash2 defines a hyperparameter set for the R10.4.1 pore version. For these reasons, we compare Campolina segmentation with two versions of the Scrappie algorithm for the R9.4.1 datasets, and one version of the Scrappie algorithm for the R10.4.1 datasets. The sets of Scrappie hyperparameters defined by Sigmoni and RawHash2 are available in the Supplementary Material S3.

The segmentation evaluation results are shown in Figure 4(b). Campolina segmentation proves to be of better quality than Scrappie for all datasets, across different pore versions, and based on a wide range of evaluation metrics. Campolina identifies more true border positions, based on Jaccard similarities, and is generally closer to actual border positions, based on Chamfer distance. Based on Alignment ratios and Alignment score, the obtained segmentation aligns well with the ground-truth segmentation. Finally, the z-normalized mean values of the obtained events correspond well to the aligned ground-truth events, based on the L1 distance, and to the expected levels on the reference, based on Pearson’s correlation coefficient *r*.

Campolina’s segmentation reduces the over-segmentation errors present in Scrappie while also reducing the skip errors where an event border is not detected. Furthermore, the event levels identified by following Campolina’s segmentation align more consistently with the ground truth, resulting in longer, uninterrupted chains of correctly segmented events. The continuity reflects accurate detection of event levels that correspond closely to the ground truth levels and the corresponding k-mer levels. These improvements are quantitatively supported by a significantly lower L1 distance to the ground truth event means and a substantially higher Pearson’s correlation *r* with the expected reference levels.

The foundation of the real-time approaches is a successful matching of the segmented event levels in the signals with the k-mer levels on the reference. Given the high signal to noise ratio in raw nanopore signals and a narrow distribution of possible k-mer levels, a segmentation that provides events that correspond to the given reference tightly, such as demonstrated by Campolina, is crucial for developing raw signal processing frameworks that can accurately and efficiently match raw nanopore signals to the reference. By generating event levels that match the reference with high fidelity, Campolina’s segmentation enables downstream processing algorithms to operate with greater accuracy and efficiency. The improved correspondence between segmented events and k-mer levels not only improves immediate alignment accuracy but also enhances the reliability and speed of real-time analyses, which is critical in time-sensitive applications.

### Campolina is well-suited for the existing raw signal processing frameworks without any adjustments to the pipelines

To further assess the quality and suitability of Campolina segmentation, we run a set of classification experiments with the existing raw signal processing frameworks, Sigmoni and RawHash2, which rely on signal segmentation, and test the effects of replacing their original segmentation with the segmentation predicted by Campolina. In this way, we assess whether Campolina’s segmentation is well-suited for the existing raw-signal processing frameworks, beyond assessing the quality of the obtained segmentation with respect to ground truth. We choose binary and multi-class classification tasks since these are common use cases for such tools. In the original work, Sigmoni was primarily evaluated on the R9.4.1 nanopore version, but we extended these experiments to include the R10.4.1 datasets. For evaluation, we randomly select 1,000 reads from each species in the D6322 dataset and 8,000 reads from the human dataset. To assign ground truth labels to the D6322 reads, we use minimap2 to map them to reference genomes. Only reads that uniquely map to a single reference and with a mapping quality of 60 are assigned the corresponding label.

For binary classification, we utilize the D6322 dataset to distinguish between yeast and bacterial reads. This experiment is referred to as “Zymo Binary Classification”. Furthermore, we use both human and D6322 datasets to differentiate between human and bacterial reads. This experiment is referred to as “Host depletion”. As a multiclass classification task, we define “Zymo multiclass classification” where we distinguish between seven microbial species: *S. aureus, S. enterica, P. aeruginosa, L. monocytogenes, E. faecalis, B. subtilis*, and *S. cerevisiae*.

In our evaluation, the parameters and the processing steps of the original Sigmoni and RawHash2 frameworks are not modified apart from replacing the signal segmentation. We repeat each run of the algorithm twice: once without any alterations and again with the Scrappie segmentation replaced by the Campolina predicted borders. We refer to the original algorithm used in Sigmoni or RawHash2 as Scrappie, although the segmentation hyperparameters used in their implementations vary. All Scrappie hyperparameters were kept as defined in the frameworks. Since Sigmoni does not provide a separate set of R10.4.1 Scrappie hyperparameters, we run the framework with the default one because that hyperparameter setup achieves the best results for Scrappie segmentation. The Sigmoni and RawHash2 commands we ran for different classification experiments are available in the Supplementary Material S1.

While the Sigmoni framework assigns a label to each input example, the RawHash2 framework labels an example as “unclassified” when it cannot be mapped to any provided reference. In our evaluation, we treat the “unclassified” examples as misclassified when calculating classification metrics, but provide details on the “unclassified rate” that show the percentage of the examples assigned the “unclassified” label.

We report read length-weighted accuracy, precision, recall and F1 score for R10.4.1 datasets and R9.4.1 datasets.

Although Sigmoni and RawHash2 were originally developed and optimized for the Scrappie segmentation and the R9.4.1 nanopore version, our segmentation method, Campolina, demonstrates consistently superior or comparable performance across all classification tasks and both nanopore versions (R10.4.1 and R9.4.1), as shown in Figure 5. We provide detailed per-class metrics, precision, recall, and F1, true positives, false positives, false negatives, and unclassified rates for both frameworks and both segmentation tools in the Supplementary Tables 3-8 and Supplementary Material S5.

**Figure 5.**
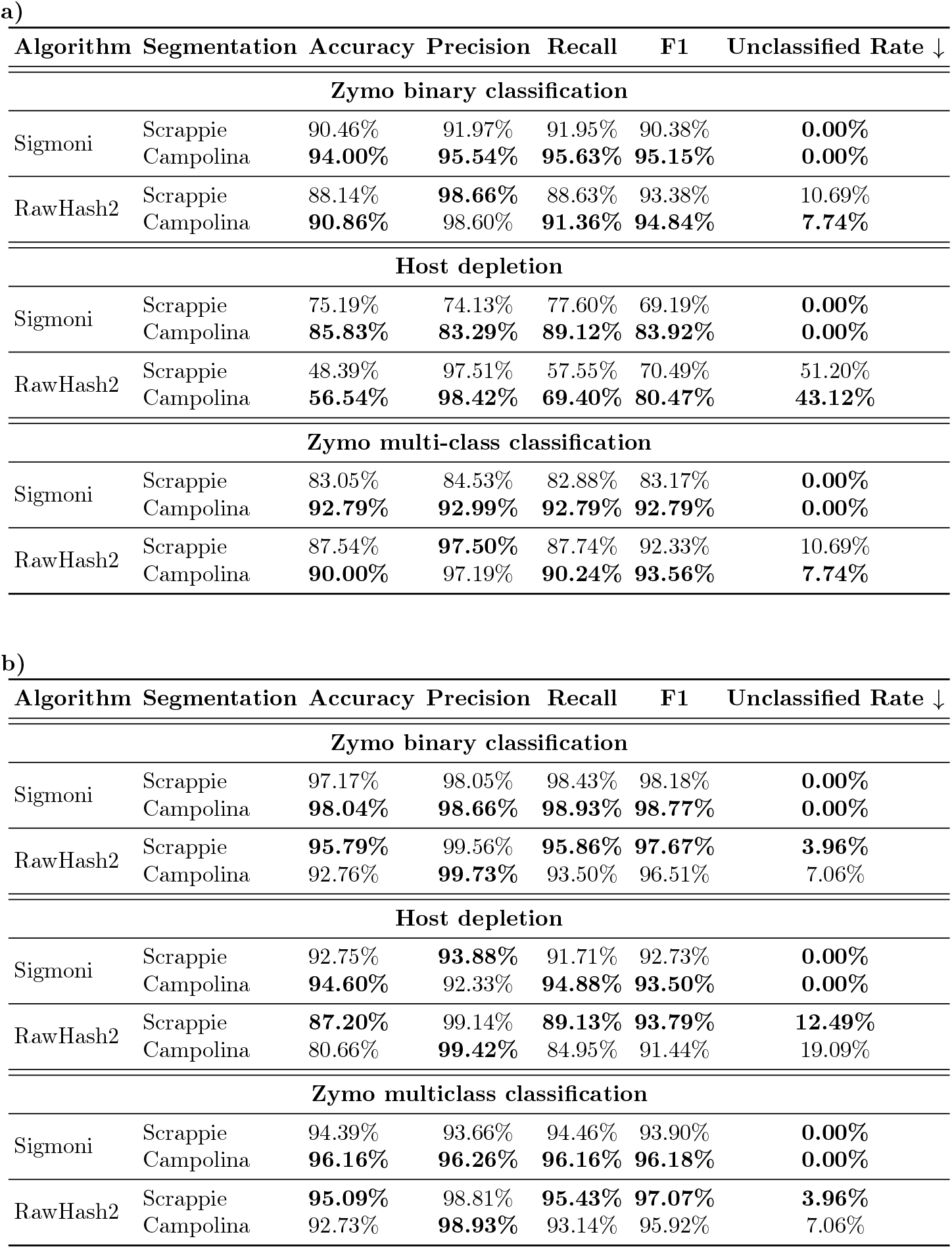
Segmentation suitability evaluation. **a)** Results for the R10.4.1 nanopore version. **b)** Results for the R9.4.1 nanopore version. Performance is evaluated on three tasks: Zymo binary classification, Host depletion, and Zymo multiclass classification. Metrics reported include length-weighted accuracy, precision, recall, F1 score, and unclassified rate. Unclassified rate is the percentage of reads that could not be assigned to any of the provided references in the RawHash2 framework. We treat those reads as misclassified when calculating other metrics, but also provide the details on the “unclassified rate” to quantify the ratio of reads that cannot be aligned to any reference. Both segmentation strategies (Scrappie and Campolina) are evaluated using two classification frameworks (Sigmoni and RawHash2) and three classification tasks.

On the R10.4.1 nanopore version, as shown in Figure 5(a), Campolina yields substantial gains in all tasks and for both raw signal processing frameworks. The unclassified rates are reduced with increased accuracy in the RawHash2 framework, and classification accuracy is significantly improved compared with the default Sigmoni framework.

Campolina consistently improves Sigmoni on the R9.4.1 nanopore version as well (Figure 5(b)), but slightly underperforms compared with Scrappie in the RawHash2 framework, mainly due to slightly higher unclassified rates.

It is important to highlight that both frameworks were originally developed and optimized for Scrappie segmentation, including formulating specially tailored postprocessing steps that aim to resolve segmentation errors, and the R9.4.1 nanopore version. Therefore, the results Campolina achieves across different tasks, frameworks, and nanopore versions prove the robustness, adaptability, and effectiveness of Campolina as a general-purpose segmentation strategy. We hypothesize that the results achieved in the existing frameworks with Campolina as a segmentation choice could be further improved by making adjustments to the segmentation post-processing scheme, but deem such experiments out of scope.

### Campolina is time efficient and appropriate for real-time frameworks

In addition to the high quality of the segmentation, an important aspect of a segmentation framework is the execution time. To assess the resource utilization of different approaches, we utilize 8000 reads from the R10.4.1 NA12878 dataset.

We compare the resource requirements of Campolina frameworks in two ways. First, we assess the running time along with CPU and memory usage for the full Campolina framework and the ground truth pipeline. We run each setup five times and report the mean execution time in seconds. The comparison of the time required to obtain precise event borders through refinement and the time required to obtain a segmentation of comparable quality with Campolina, which does not depend on the basecalling, alignment, and refinement steps, is shown in Figure 6(a).

**Figure 6.**
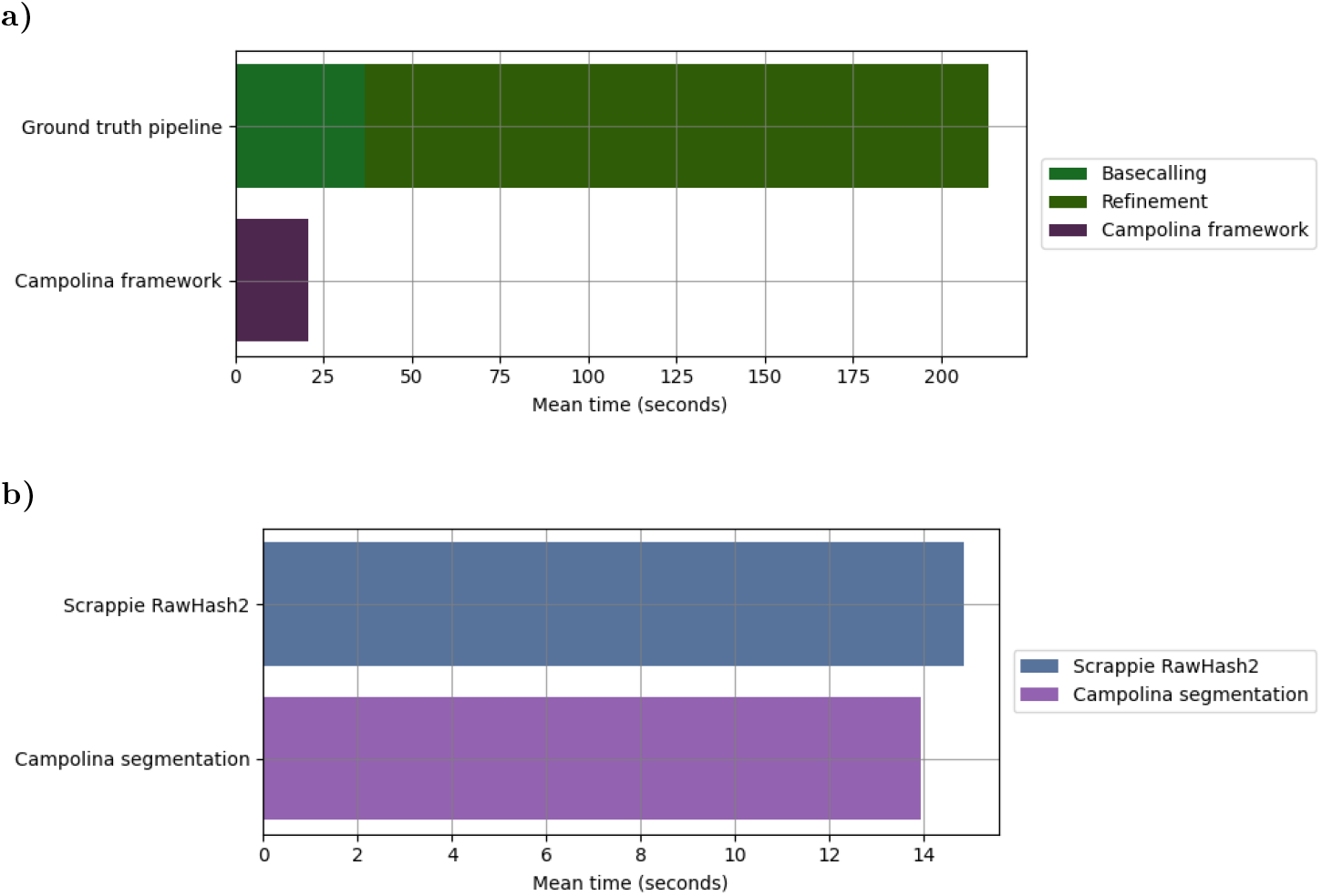
Time requirement analysis. **a)** Comparison of total resource usage and execution time (mean of 5 runs) between the full Campolina framework and the ground truth pipeline (high-accuracy basecalling + refinement). **b)** Comparison of segmentation-only execution time (mean of 5 runs) between Campolina and Scrappie. Campolina performs augmentation, deep neural network processing, and border inference on a GPU, while Scrappie runs on a singlethreaded CPU setup.

The basecalling step in the ground truth is executed on a single A100 GPU, while the refinement is run on CPU in a single-thread setup. Campolina’s inference is executed on a single A100 GPU, while the storing of the border positions in Parquet format is executed in a separate CPU thread. Second, we compare the time required for obtaining border positions by Campolina and Scrappie. We do so by only measuring the time that the Campolina framework spends on segmentation-related tasks: augmenting raw signal chunks with statistical descriptors, processing the chunks by the deep neural network, and inferring border positions in the provided chunks based on the model output. Similarly, we measure the time Scrappie’s algorithm requires to process signal chunks and return event border positions. We measure the execution time five times for both frameworks and report the mean execution time in seconds. The execution time comparison is shown in Figure 6(b). The Scrappie algorithm is executed solely on the CPU and in a single-thread setting. We acknowledge the differences in the resource requirements of Campolina and Scrappie. Campolina, being a deep neural network, is primarily designed for execution on a GPU, while Scrappie runs completely on a CPU.

We do not compare Campolina segmentation with the Scrappie implementation used in Sigmoni for technical reasons. Sigmoni is a tool primarily implemented in Python, but utilizes the Scrappie algorithm provided as part of the Uncalled4 framework ^3^. Uncalled implements Scrappie in C++ (equivalent to RawHash2 implementation) with a Python binding that allows Sigmoni to provide a Python array to obtain the normalized means of the predicted events. Given that Sigmoni invokes a function that includes both extracting the border positions and calculating the normalized means of the obtained events, and that the provided C++ Scrappie implementation corresponds completely to the RawHash2 implementation with the only difference being the computational overhead coming from bridging Python and C++, we do not make a direct comparison to Scrappie implemented in Sigmoni tool. However, we provide a comparison of Scrappie segmentation + event mean extraction execution time in RawHash2 and Sigmoni in the Supplementary Material S4.

The analyses in Figure 6 show that direct segmentation done by Campolina significantly reduces the time required for obtaining accurate signal borders when compared with the reference-guided basecalling and refinement approach. Additionally, when compared with the traditional algorithmic segmentation done by Scrappie, under the assumption of different resource requirements, Campolina is well-suited for real-time raw signal processing frameworks with an execution time that is comparable to Scrappie’s execution time. Furthermore, since Campolina is adapted to processing chunks of signals rather than full signals and can operate on batches of chunks simultaneously, it is an appropriate segmentation framework for the existing raw signal processing frameworks, such as RawHash2 and Sigmoni.

Comprehensive information on resource utilization is available in Supplementary Table 1 and Supplementary Table 2.

## 3 Discussion

We present Campolina, a first deep-learning framework for high-quality segmentation of raw nanopore signals. Campolina is trained separately for the R9.4.1 and R10.4.1 nanopore versions and thoroughly evaluated against the traditional algorithmic approach, Scrappie. The focus of this manuscript is on the R10.4.1 version of nanopores. However, we also included R9.4.1 version in our analysis despite its discontinued official support by ONT, given the substantial volume of existing data generated with this version, and to demonstrate the consistency of our method across different nanopore versions. Campolina is based on a convolutional architecture and trained in a supervised manner using refined event borders from the Dorado-Remora framework as ground truth. This ground-truth pipeline aligns with the core components of ONT’s nanopore sequencing development, incorporates the approximation of segmentation from the latest basecallers, via a move table, alignment to the reference, and access to nanopore characteristics, i.e., k-mer models, in combination with a dynamic programming strategy to find the best border positions in the given signal. Campolina is optimized with a composite loss: Focal loss for classification accuracy, Huber loss to constrain the number of predicted events, and a custom Consecutive loss to prevent redundant border predictions. To assess the quality and suitability of Campolina segmentation, we perform a two-step evaluation. We present Campolina, a first deep-learning framework for high-quality segmentation of raw nanopore signals. Campolina is trained separately for the R9.4.1 and R10.4.1 nanopore versions and thoroughly evaluated against the traditional algorithmic approach, Scrappie. The focus of this manuscript is on the R10.4.1 version of nanopores. However, we also included R9.4.1 version in our analysis despite its discontinued official support by ONT, given the substantial volume of existing data generated with this version, and to demonstrate the consistency of our method across different nanopore versions. Campolina is based on a convolutional architecture and trained in a supervised manner using refined event borders from the Dorado-Remora framework as ground truth. This ground-truth pipeline aligns with the core components of ONT’s nanopore sequencing development, incorporates the approximation of segmentation from the latest basecallers, via a move table, alignment to the reference, and access to nanopore characteristics, i.e., k-mer models, in combination with a dynamic programming strategy to find the best border positions in the given signal. Campolina is optimized with a composite loss: Focal loss for classification accuracy, Huber loss to constrain the number of predicted events, and a custom Consecutive loss to prevent redundant border predictions. To assess the quality and suitability of Campolina segmentation, we perform a two-step evaluation.

First, we measure the quality of obtained segmentation through a carefully designed evaluation pipeline that exceeds naive border position matching evaluation and ensures a fair and complete assessment of segmentation quality. The proposed segmentation quality assessment quantifies how well the predicted borders match the ground truth ones, but also measures the quality of the obtained events by comparing event levels with the corresponding ground truth events, and the expected k-mer levels. Second, we explore the suitability of Campolina segmentation in the existing raw-signal processing frameworks, Sigmoni and RawHash2, by replacing the traditional segmentation with borders predicted by Campolina, and running the frameworks on several downstream tasks.

Campolina improves segmentation quality over Scrappie in several key aspects. The predicted border positions match the ground truth more precisely, and the alignment between the predicted and the ground truth sequence of events is better. Additionally, Campolina significantly reduces both oversegmentation errors, where events are unnecessarily split, and stay errors, where signal changes are missed. These improvements are particularly important for raw-signal processing frameworks that rely on precise event segmentation. Moreover, Campolina not only matches the ground-truth segmentation more closely than Scrappie but also aligns better with the expected k-mer levels. Existing raw-signal processing frameworks often apply extensive post-processing steps to correct segmentation errors introduced by Scrappie. These include custom heuristics and additional filtering aimed at mitigating issues such as oversegmentation. The need for such substantial post hoc correction highlights a fundamental limitation in the initial segmentation quality provided by Scrappie. Additionally, the existing segmentation corrections largely overlook the stay errors, where a meaningful event is missed entirely, and no border is placed where one should be. Our perturbation analysis reveals that deletions of the borders have a greater impact on downstream performance than oversegmentation, yet they remain unaddressed. Campolina simultaneously reduces the insertion and deletion errors, reduces the need for corrective post-processing, and provides cleaner, more reliable input for subsequent analysis.

Moreover, when selected as a segmentation strategy in the existing raw signal processing frameworks, Sigmoni and RawHash2, Campolina improves the classification accuracy across different tasks and for both frameworks in the R10.4.1 datasets. Even for the R9.4.1 nanopore version, for which the frameworks were initially developed, Campolina improves the Sigmoni framework, and performs comparably to RawHash2 with a slightly higher unclassified rate. These results are achieved without any alterations to the post-segmentation pipeline despite being developed for Scrappie segmentation. Given that the quality of Campolina segmentation is significantly higher compared with Scrappie segmentation, we hypothesize that some adjustments to the existing raw signal processing frameworks, especially related to correction of segmentation errors, could, with integrated Campolina segmentation, improve the overall performance of the existing tools, and open possibilities for further development of real-time frameworks.

One of the limitations of the framework we propose is the dependency on a two-step pipeline for obtaining the ground-truth border positions. However, we deem this approach well aligned to the core development of nanopore sequencing technology and argue that the ground truth pipeline that we rely on could be replicated as long as there is a way to extract initial approximation of the segmentation, align the basecalled signal to the reference and describe the k-mer level distribution for a specific nanopore version. Campolina is a deep-learning-based framework, and despite being lightweight for efficiency, it requires more computational resources and is primarily designed for processing on a GPU compared with traditional algorithmic segmentation that runs on CPU. However, with the quality of the obtained segmentation being significantly higher, and the Campolina framework being designed for a fast execution on a single GPU, we argue that the resource requirement trade-off is reasonable. The resource utilization analysis we perform show a clear advantage of direct segmentation to the reference-guided pipeline that we utilize for obtaining ground truth. Even when compared with Scrappie’s execution time, Campolina proves as an appropriate framework for real-time processing frameworks. Additionally, we hypothesize that the increased segmentation quality could impose simplifications or removal of some post-segmentation processing steps, therefore introducing time savings. In future work, we plan to explore the suitability of the Campolina framework for the segmentation of RNA signals, as well as optimize, further evaluate, and integrate Campolina segmentation into the existing frameworks for processing raw nanopore signals. Currently, Campolina is implemented entirely in Python and would benefit from optimization to improve computational efficiency; however, we consider such experiments out of the current scope.

## 4 Conclusions

This study presents Campolina, a lightweight deep learning-based framework for the segmentation of raw nanopore signals, which significantly outperforms the traditional algorithmic approach, Scrappie. Campolina delivers more accurate and consistent segmentation across different nanopore versions (R9.4.1 and R10.4.1) and species, producing event sequences that better align with ground truth borders and expected k-mer signal levels. These improvements reduce both oversegmentation and stay errors, which are critical for the accuracy and efficiency of downstream raw signal processing.

Importantly, Campolina enhances the performance of existing frameworks such as Sigmoni and RawHash2 without requiring any changes to their post-segmentation pipelines. This demonstrates the practical relevance and compatibility of the proposed method within current signal processing ecosystems. Despite requiring GPU resources, Campolina offers real-time performance and introduces the possibility of simplifying or removing error-correction steps commonly needed with Scrappie.

The improved segmentation quality provided by Campolina opens the door for developing faster, alignment-free classification methods based on simple hash matching, skipping the alignment step proposed in RawHash2 or the r-index building proposed in Sigmoni, which could redefine how raw nanopore signals are processed. As such, we consider high-quality segmentation an important step toward more accurate, scalable, and real-time nanopore signal analysis frameworks.

## 5 Methods

### 5.1 Ground Truth Pipeline

To train Campolina models, we first obtain the ground-truth positions of event borders in the signals. These positions are used as labels in the supervised border detection learning pipeline. To get the ground-truth event borders for our datasets, we follow a two-step pipeline where we basecall and align the signals with Dorado and then refine the signal mapping with the Remora suite.

In the first step, raw nanopore signals are basecalled with the “super accurate” Dorado model (version 0.5.1 for R9.4.1 data, version 0.4.2 for R10.4.1 data) to ensure high basecalling quality. In the basecalling step, we provide the reference, which is used to get the alignment of the read to the reference. Additionally, we utilize Dorado’s option to extract the move table. The move table approximates the signal-to-sequence alignment and serves as a starting point for obtaining accurate event borders. Both the alignment, and the move table are used in the refinement step to adjust the signal scaling and the signal-to-sequence mapping to more closely match the expected signal levels.

In the second step of the ground truth pipeline, we rely on a signal-to-sequence mapping refinement algorithm provided as an auxiliary option integrated into the Remora suite. Remora’s refinement is based on a dynamic programming step that takes the input signal-to-sequence alignment defined with the move table and searches for the optimal mapping in a band that extends bases on either side. The refinement can be anchored on the basecalled sequence or the reference mapping. We perform the refinement anchored on the reference mapping to mitigate basecalling error effects and get the most accurate signal segmentation.

Remora adjusts the signal-to-sequence mapping based on the expected event levels that are extracted from a k-mer model. The k-mer model is a lookup table that defines the expected values for every possible k-mer in the pore. The lengths of the k-mers in the pore are defined by the pore version. K-mer length for the R9.4.1 pore is 6, while the R10.4.1 pore version has a k-mer of length 9. Oxford Nanopore Technologies (ONT) provides k-mer models for both pore versions, but with differences in formatting. The R9.4.1 k-mer model defines the mean and standard deviation of a Gaussian distribution representing the current observed for a specific k-mer with additional information on mean, standard deviation, and lambda parameter of the inverse Gaussian distribution modeling corresponding signal noise. The R10.4.1 k-mer model only provides the expected mean value for each k-mer and z-normalizes the values across the full lookup table. Since we rely on the Remora suite for refinement, and the refinement is developed for the R10.4.1 k-mer model format, we extract and z-normalize the R9.4.1 mean levels from the original k-mer model and use them in the refinement step in an appropriate format. We provide Dorado and Remora commands to extract ground truth border positions in the Supplementary Material S1.

### 5.2 Feature Extraction

The preprocessing of raw signals includes splitting the signals into chunks of fixed length and a simple channel augmentation. A simple channel augmentation has proven to be effective in enhancing the performance of convolutional networks in a wide range of applications [20]. In the first step, we split the raw signals into chunks of uniform length. The chunks are nonoverlapping, with chunk length set to *L*=6000 samples in all experiments. The final chunk of the signal is padded with zeros, if needed, to ensure the target length. Each chunk of the signal is z-normalized to mitigate potential batch effects. The raw signal values are augmented with four additional descriptors, *µ, σ, diff1, t-statistics* as shown in Figure 1(a).

First, we slide across the signal and calculate the mean (*µ*) and standard deviation (*σ*) for rolling windows of length *w*_*m*_ = 3. Furthermore, we compare the consecutive signal values and calculate the change in signal values between two consecutive points (*diff1*). Finally, we calculate *t-statistics* for rolling windows of length *w*_*t*_ = 3 following Welch’s t-test formulation [21].

Each input example is obtained by appending signal descriptors to the z-normalized signal chunk, which results in a 6000 *×* 5 dimensional input.

### 5.3 Architecture

Campolina’s architecture consists of two main components: a convolutional network and a classification head as shown in Figure 1(b).

The convolutional network is built of sequential convolutional blocks, where each block contains a convolutional layer followed by a non-linear activation and a batch normalization. The convolutional component of our models consists of five convolutional blocks with output channel dimensions set to 32, 64, 64, 128, 128, in that order. Kernel size is set to 3 in the first convolutional block, and to 31 in all subsequent blocks. The activation function is GELU in all convolutional blocks.

The classification head is a single linear layer that outputs a non-normalized event border probability for each point in the input example.

The final output of the model is of shape *L ×* 1 with a single non-normalized probability prediction for each input point being a border.

The architecture setup explained above was used for both the R9.4.1 and R10.4.1 versions of Campolina. The architectures choices were made to keep the model lightweight because of time and computational requirements, but capable of modeling relevant local interactions. The kernel of size 3 in the first layer gathers short signal segments that potentially contain an event-change, while the subsequent layers with a wider kernel increase the receptive field to roughly capture all events of the signal that include a certain nucleotide in the sequence.

### 5.4 Training and Evaluation

Campolina is trained end-to-end in a supervised manner with the objective of correctly identifying the event borders.

#### 5.4.1 Border Detection Task

We model the border detection problem as a point-wise binary classification task where each input point is classified as a border or not a border. Event border indices obtained as groundtruth are expanded into binary vectors of signal chunk lengths, where each 1 in the vector indicates that a specific input point is a border. Given that the ratio between the number of border and non-border points is extremely imbalanced, we choose Focal loss as the classification loss. Focal loss is an adaptation to the binary cross-entropy loss (BCE) designed to address the extreme imbalance in datasets. Having an estimated probability *p ∈* [0, 1] for class *y* = 1, we define

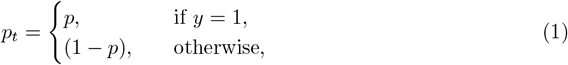

and formulate BCE= −log(*p*_*t*_). To address the imbalance issue, BCE is adapted to Focal loss with two additional parameters, *α* and *γ. α* is a weighting factor that accounts for imbalance by assigning higher values to the sparse class. We define *α*_*t*_ following the principle from Equation

1. In addition, Focal loss addresses easily classified negatives that comprise the majority of the loss and dominate the gradient with a modulating factor *γ*. The modulating factor reduces the loss contribution from easy examples and extends the range in which an example receives a low loss. The final formulation of the Focal loss FL is

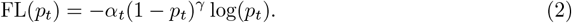

We set *α*=0.8 and *γ*=1 in all our experiments.

#### 5.4.2 Auxiliary Losses

We introduce two auxiliary losses during the training to improve learning and mitigate problems related to the variable speed of DNA translocation through a pore.

##### Huber Loss

Because of a poor signal-to-noise ratio, over-segmentation of nanopore signals is a common problem. To prevent the model from over-segmenting the signals, we introduce the Huber loss [22] component that constrains the number of predicted borders by comparing the predicted probability vector and the label vector. Huber loss is a quadratic function for absolute error values that fall below the predefined *δ*, and *δ*-scaled L1 loss otherwise. For an *L*-dimensional vector of normalized point-wise predictions **p**, *∀* i=1, …, *L p_i_* ∈ [0, 1], and a label

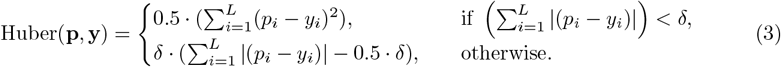

The Huber threshold *δ* = 20 in our experiments.

##### Consecutive loss

The non-constant translocation speed of DNA through the nanopore affects the distribution of event lengths and the number of samples measured as the nucleotide in the nanopore changes. We often observe several consecutive samples that correspond to measurements taken during the event change. To prevent the model from predicting multiple consecutive points as event borders, we introduce the Consecutive loss. To calculate the Consecutive loss component, we start by expanding the *L*-dimensional vector of normalized probabilities **p** to two (*L* + 1)-dimensional vectors, **p**00 with a zero inserted at position 0 and **p**_*L*0_with a zero inserted at the end of the vector **p**. The consecutive loss of the vector **p** is calculated as follows:

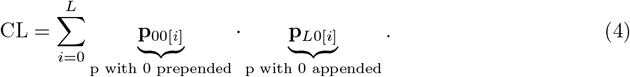

This way, we penalize giving a high score to two consecutive points with a high product between the **p**_00_ and **p**_*L*0_vectors.

#### 5.4.3 Final Loss and Optimization

The final loss for the training is given as a linear combination of three loss components:

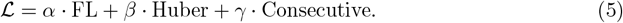

*α, β,* and *γ* account for the different scales of loss components and are set to *α*=5000, *β*=0.05, *γ*=0.1 in our experiments.

All model weights are optimized using a modified version of Adam [23], which decouples weight decay from the optimization procedure [24]. The learning rate is set to 1*e*^−3^ in our experiments. Running average coefficients *β*_1_, *β*_2_ and weight decay are left at their default values *β*_1_ = 0.9, *β*_2_ = 0.999, weight decay = 1*e*^−2^.

### 5.5 Datasets

Segmentation quality and suitability are evaluated using two datasets sequenced with R9.4.1 and R10.4.1 nanopore version.

The R9.4.1 Zymo D6322 dataset is utilized for quantifying the segmentation quality and in all three classification tasks. The species included in this analysis are: *S. aureus, S. enterica, P. aeruginosa, L. monocytogenes, E. faecalis, B. subtilis*, and *S. cerevisiae*. We randomly select 1,000 reads from each Zymo species and assign ground truth labels only to reads that map uniquely to a single reference with a mapping quality of 60. *E. coli* reads from the dataset were used for training and excluded from evaluation.

Human dataset used for R9.4.1 evaluations is the B-lymphocyte cell line: NA12878 [25]. 8000 reads are randomly sampled from the original dataset for evaluation.

*E. coli* reads from the R10.4.1 Zymo dataset were used for training. The species included in the evaluations are: *S. aureus, S. enterica, P. aeruginosa, L. monocytogenes, E. faecalis, B. subtilis*, and *S. cerevisiae*.

The R10.4.1 human evaluation dataset was the NA12878 dataset from which we randomly sampled 8000 reads for evaluation purposes.

#### 5.5.1 Evaluation

We perform a two-level evaluation of the obtained segmentation, one that assesses the quality of obtained events with respect to the ground truth segmentation, and expected event levels given the corresponding reference sequence and alignment information, and the other that evaluates the suitability of the Campolina segmentation in the existing frameworks that rely on the segmentation algorithms.

##### Segmentation Evaluation

Evaluation of nanopore signal segmentation is a non-trivial problem. While the naive approach to the evaluation would be to compare the sets of predicted and true borders, we argue that this information is insufficient for a truthful assessment of segmentation quality. Since it is common to observe multiple measures taken as the nucleotide in the pore changes, the choice for the event-change point in the signal can be nonunique. The experiments with border perturbations presented support for our hypothesis that local border perturbations do not have a strong effect on segmentation quality.

To ensure a fair evaluation of the segmentation and overcome the previously mentioned challenges, we outline a set of metrics that jointly assess the quality of nanopore signal segmentation. The predicted segmentation is compared with the positions obtained through the ground truth pipeline. Additionally, the obtained events are compared with the expected values defined by the reference sequence and mapping of the signal to the reference. We evaluate the quality of segmentation obtained by Campolina and Scrappie.

We start by comparing the sets of borders predicted by Campolina or Scrappie and borders obtained as ground truth and calculating the Jaccard similarity of the two sets. If we have a set of predicted border positions A, and a corresponding set of ground truth border positions B, we calculate the *Jaccard similarity (J)* as

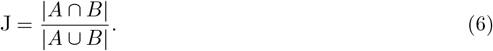

Additionally, we define the expanded Jaccard similarity where we look for the intersection between two sets by comparing predicted and labeled positions while taking into account their ±1 surrounding. Having a prediction position *b*_*A*_, and a labeled position *b*_*B*_, the border pair is counted at the intersection of two sets if *b*_*A*_ = *b*_*B*_ *−* 1, *b*_*A*_ = *b*_*B*_, or *b*_*A*_ = *b*_*B*_ + 1.

Furthermore, we assess the quality of signal segmentation by calculating a bi-directional *L1 Chamfer distance (CD)* - a common metric in point cloud tasks - for the set of predicted borders A and the set of ground-truth borders B capturing how well they align by measuring distances between the closest pairs as follows:

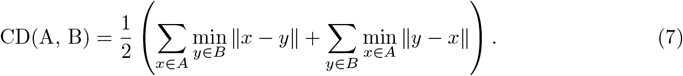

Although the expanded Jaccard similarity accounts for some cases of local border position perturbations, and bi-directional Chamfer distance gives insight into how well the predicted borders align with the ground truth, we proceed by aligning Campolina segmentation to ground truth segmentation arguing that good alignment between the obtained segmentation and the groundtruth segmentation implies that the predicted events are of high value for various downstream tasks. The alignment process goes as follows: we index the signal, group indices into events based on event border positions, and form two sets of index subsets. We compare the index subsets and compute a *Match Score (MS)* for each predicted–ground-truth pair based on index overlap:

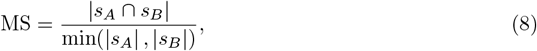

where *s*_*A*_ and *s*_*B*_ are the sets of ground truth and predicted events, respectively. MS quantifies how well a signal stretch defined by the ground truth segmentation matches each predicted event with the goal of finding one event that corresponds to the ground truth event best. MS *∈* [0, 1]. Having calculated the match scores, we construct a *N*_*A*_ × *N*_*B*_ matrix where *N*_*A*_ is the number of ground-truth events, and *N*_*B*_ is the number of predicted events. Starting from the last event pair with a match score larger than zero, we initiate the traceback - the reconstruction of the optimal alignment by following the path of the highest scores in the alignment matrix starting from the final aligned pair. The alignment matrix is indexed from the upper left to the lower right corner, where we initiate the traceback, and in each step of the traceback process, we move by one position to the left, up, or diagonally left-up. The move is decided based on the MS values in the neighboring cells. From the cell in row *i* and column *j* we move as follows

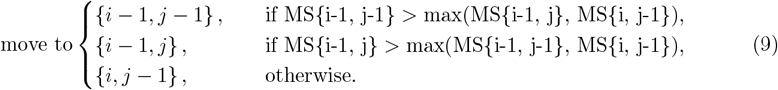

After obtaining full alignment between predicted segmentation and ground-truth segmentation, we proceed by defining three possible relations between aligned events:

- match (m) = one predicted event is aligned to one ground-truth event
- insertion (i) = multiple predicted events are aligned to one ground-truth event
- deletion (d) = one predicted event is aligned to multiple ground-truth events

Having a sequence *A* of event alignments with each alignment *a* = *m, i, d*, we measure the quality of alignment by calculating the *Alignment Score (AS)* as follows

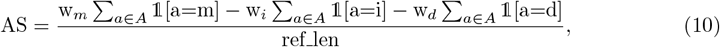

with score weights set at w_*m*_ = 1, w_*i*_ = 0.5, w_*d*_ = 0.5 and ref len defined as the overall number of ground-truth events that we aligned to the predicted events. This scoring penalizes frequent short insertions more than fewer longer insertions. We adopted this formulation since the continuity in accurate segmentation is important in existing pipelines that rely on consecutive events for further processing, i.e., RawHash2 chains consecutive events for further processing, and a larger number of insertion blocks interrupts the continuity more.

Additionally, we find the ratio of the ground truth events that are in match, insertion and deletion relation with Campolina events and report those values.

Finally, based on the alignment between two sets of events, and alignment of ground-truth events to the reference (obtained through basecalling step of the ground-truth pipeline by providing the reference as an input argument), we anchor the predicted segmentation to both ground-truth segmentation and the corresponding reference sequence and evaluate the quality of the obtained segmentation in the following ways:

1. We calculate the *L1 difference* between z-normalized event means for the aligned segmented- ground-truth event pairs.
2. We calculate *Pearson correlation coefficient r* between z-normalized segmented events and the expected event level as defined in the appropriate k-mer model for the k-mer corresponding to the segmented event based on the segmented event-ground-truth and ground-truth-reference alignments.

The absolute difference between the normalized values of the predicted event and the aligned ground-truth event gives insight into how well the segmentation matches the expected normalized values. Similarly, calculating Pearson’s *r* between the predicted segmentation and the corresponding k-mer model values quantifies how well the segmented signal matches the reference sequence. The values extracted from the k-mer model are z-normalized with respect to the whole k-mer model, while the predicted events are normalized with respect to the signal. We choose Pearson’s *r* instead of absolute difference to mitigate the potential bias from normalizing on a signal level compared to normalizing on the k-mer model level.

##### Ablation Study

In our experiments, we build a multi-component loss function that, in addition to classification loss measured by Focal loss, constrains the number of predicted events with Huber loss and prevents the model from predicting consecutive points with the custom Consecutive loss. To assess the contribution of each component in our training setup, we perform ablation by training two additional models, one with only the Focal loss component and the other with Focal + Huber loss components. We predict event borders on the R10 Zymo dataset and measure the segmentation quality with the proposed evaluation pipeline, as well as the suitability of the obtained segmentation in RawHash2 on “Zymo multiclass classification”. The Huber and Consecutive loss components improve the quality of the obtained segmentation which is mirrored in the increased number of matches in the alignment, reduced L1 distance of the obtained to the ground truth event levels, and increased correlation with the reference k-mer levels. Moreover, Campolina trained with a three-component loss reduces the unclassified rate and improves the classification accuracy when compared with the ablated versions on a “Zymo multiclass classification” task with RawHash2. Full ablation results are available in the Supplementary Material S2.

## 6 Declarations

### 6.1 Data availability

The Zymo D6322 dataset for the R9.4.1 nanopore version was previously sequenced by Kovaka et al. (2021) from the ZymoBIOMICS High Molecular Weight DNA Mock Microbial Community. The R9.4.1 NA12878 dataset is available via AWS at https://github.com/nanopore-wgs-consortium/NA12878. The R10.4.1 Zymo dataset was internally sequenced. The R10.4.1 human evaluation dataset was the NA12878 dataset (Oxford Nanopore Technologies Benchmark Datasets was accessed on 2024-01-02 from https://registry.opendata.aws/ont-open-data).

### 6.2 Code availability

The code repository is available at https://github.com/lbcb-sci/Campolina.git, and the pretrained R9.4.1 and R10.4.1 models are available at https://zenodo.org/records/15626806. The weights can be easily downloaded using the script provided in the repository. We provide scripts and instructions for training, inference, and segmentation evaluation in the code repository.

### 6.5 Author contributions

S.B. designed and implemented Campolina with help from K.F. and M.Š.; S.B. and K.F. performed bioinformatics analysis and evaluation with the contribution of M.Š.; S.B., K.F., and M.Š. organized the manuscript. S.B., K.F., B.H. and M.Š. wrote the manuscript. M.Š. supervised the project. M.Š. and B.H. provided mentorship and support during the project.

## Footnotes

1 https://nanoporetech.com/products/sequence/minion

2 https://nanoporetech.com/products/sequence/gridion

3 https://github.com/skovaka/uncalled4/blob/main/src/cpp/eventdetector.cppL249

