## Supplementary information for "Campolina: A Deep Neural Framework for Accurate Segmentation of Nanopore Signals"

#### S1 Tool versions and commands

**Dorado basecalling R9.4.1** The R9.4.1 datasets are basecalled using the "super accurate" Dorado basecaller dna_r9.4.1_e8_sup@v3.6. The Dorado version used for basecalling is 0.5.1. Reference is provided to obtain the mapping information, and the move table is extracted by providing "–emit-moves" argument.

```
dorado basecaller -x <GPU> --emit-moves \
--reference reference.fasta -o basecalls.bam \
models/dna_r9.4.1_e8_sup@v3.6/ <reads.pod5>
```

**Dorado R10.4.1** The R10.4.1 datasets are basecalled using the "super accurate" Dorado basecaller dna_r10.4.1_e8.2_400bps_sup@v4.2.0. The Dorado version used for basecalling is 0.4.2. Reference is provided to obtain the mapping information, and the move table is extracted by providing "–emit-moves" argument.

```
dorado basecaller -x <GPU> --emit-moves \
--reference reference.fasta -o basecalls.bam \
models/dna_r10.4.1_e8.2_400bps_sup@v4.2.0/ reads.pod5
```

**Signal refinement - Remora - version 3.2.0** The signal-to-sequence mapping refinement is done using Remora's auxiliary refinement option as described in Remora Signal Mapping Refinement. The refiner is initialized with the appropriate k-mer model as follows:

```
sig_map_refiner = refine_signal_map.SigMapRefiner(
kmer_model_filename=<kmer_model>,
do_rough_rescale=True,
scale_iters=0,
do_fix_guage=True,
)
```

The refined event borders are extracted by anchoring the read (io_read) to the reference with:

```
io_read.set_refine_signal_mapping(sig_map_refiner, ref_mapping=True)
```

The extracted, refined event borders are written in the existing BAM file as a new tag labeled as "RR" and used for ground truth during Campolina training.

**Sigmoni - host depletion** To run the host depletion task, we first index the genomes, CHM13v2 as a positive reference, and bacterial genomes as the negative ones, and shred them into 100 Kbp regions for classification. Then, we classify human and Zymo reads with thresholds set as defined in the original Sigmoni work.

```
python index.py -p <path_to_pos_class_fasta_files>/*.fasta \
-n <path_to_neg_class_fasta_files>/*.fasta  \
-b <b> --shred 100000 -o <output_dir> --ref-prefix chm13_zymo

python main.py -i <_input_dir> \
-r refs/chm13_zymo -b <b> -o <output_dir> \
--read-prefix <read_prefix> --sp --thresh 0.9999999999666667
```

**Sigmoni - Zymo multiclass classification** To run the Zymo multiclass classification, we first index the Zymo genomes, treating them as positive references, and shredding them into 100 Kbp regions for classification. Then, we classify Zymo reads in the multiclass setup as defined in the original Sigmoni work.

```
python index.py -p <path_to_pos_class_fasta_files>/*.fasta \
-b <b> --shred 100000 -o <output_dir> --ref-prefix zymo

python main.py -i <fast5_input_dir> -r refs/zymo \
-b <b> -o <output_dir> --complexity --multi
```

**Sigmoni - Zymo binary classification** To run the Zymo multiclass classification, we first index the Zymo genomes, with yeast treated as the positive class, and the bacterial genomes treated as the negative class, and shred them into 100 Kbp regions for classification. Then, we classify Zymo reads with thresholds set as defined in the original Sigmoni work.

```
python index.py -p <path_to_pos_class_fasta_files>/*.fasta \
-n <path_to_neg_class_fasta_files>/*.fasta \
-b <b> --shred 100000 -o <output_dir> \
--ref-prefix zymo

python main.py -i <fast5_input_dir> \
-r refs/zymo -b <b> -o ./ \
--complexity --thresh 1.6666666666333334
```

**RawHash2 - indexing and mapping** The RawHash2 framework was run in two steps. First, the index of the references was built. For the Zymo binary and Zymo multiclass classification tasks, the reference index was built using the Zymo references. For the host depletion task combined CHM13v2 and Zymo references are used to build the reference index. Second, we map the reads to the reference index. For the Zymo binary and Zymo multiclass classification tasks, we map Zymo reads, and for the host depletion task, we map both human and Zymo reads.

```
rawhash2 -d ref.ind -p <kmer_model> -t <t> ref.fasta

rawhash2 -t <t> ref.ind <pod5_input_dir> > mapping.paf
```

### S2 Ablation study

To quantify the contributions of the composite loss function components, we perform an ablation study where we remove auxiliary loss components and measure the segmentation quality through the proposed segmentation evaluation pipeline, and the segmentation suitability in RawHash2 framework on a Zymo multiclass classification task. For the ablation study, we train two R10.4.1 models, one where the loss function consists of the Focal loss component solely (*Campolina FL*), and one where we add the Huber loss component to the Focal loss (*Campolina FL + HL*). The ablated models' segmentation quality is estimated on the R10.4.1 Zymo dataset and compared to the original model with Focal, Huber, and Consecutive loss components (*Campolina full*).

**a)**

| **Model** | **J↑** | | **CD↓** | **Alignment ratio** | | | **AS↑** | **L1 ↓** | | **Pearson's *r* ↑** | |
|---|---|---|---|---|---|---|---|---|---|---|---|
| | Naive | Expand | | Match↑ | Insert↓ | Delete↓ | | Full | Match | Full | Match |
| Campolina FL | 0.52 | 0.58 | **5.10** | 0.62 | 0.25 | **0.14** | 0.27 | 0.12 | 0.05 | 0.95 | **0.97** |
| Campolina FL+HL | **0.55** | 0.64 | 5.28 | 0.64 | 0.09 | 0.27 | 0.41 | 0.11 | 0.05 | 0.95 | **0.97** |
| Campolina full | 0.54 | **0.65** | 5.36 | **0.67** | **0.08** | 0.25 | **0.45** | **0.09** | **0.04** | **0.96** | **0.97** |

**b)**

| **Segmentation** | **Accuracy** | **Precision** | **Recall** | **F1** | **Unclassified Rate** |
|---|---|---|---|---|---|
| Campolina FL | 89.65% | **97.37%** | 89.86% | 93.43% | 8.51% |
| Campolina FL+HL | 72.08% | 93.30% | 72.52% | 81.24% | 23.06% |
| Campolina full | **90.00%** | 97.19% | **90.24%** | **93.56%** | **7.74%** |

Figure 1: **Ablation study.** The assessment of the **a)** segmentation quality and **b)** segmentation suitability for a model optimized using only Focal loss (Campolina FL), a model with a composite Focal + Huber loss (Campolina FL+HL), and a model optimized using a full loss function with Focal, Huber, and Consecutive loss (Campolina full). Models are evaluated using R10.4.1 Zymo datasets on a Zymo multiclass classification task and within the RawHash2 framework.

The segmentation quality assessment in Figure 1a show that Campolina FL primarily over-segments, which results in a higher insert ratio, and overall lower quality of the obtained segmentation with respect to ground truth and corresponding k-mer levels. Once the Huber loss component is introduced, Campolin FL+HL reduces the number of insertions, but at the cost of a higher deletion ratio. Finally, Campolina full with all three loss components improves the segmentation quality with the highest match ratio, the lowest L1 distance to the ground truth levels, and the highest Pearson's r with the expected k-mer levels. Moreover, when evaluated in the multiclass RawHash2 classification experiment, we see in Figure 1b that Campolina full reduces the unclassified rate and improves the classification accuracy compared with the ablated models.

#### S3 Scrappie hyperparameter setup

Campolina's segmentation is compared with Scrappie directly, and in the existing raw signal processing frameworks. For the purpose of evaluating the segmentation quality, we keep all Scrappie hyperparameters as defined in the frameworks. Given that Sigmoni does not provide a separate set of R10.4.1 hyperparameters, we test the framework on R10.4.1 datasets using the R10.4.1 Scrappie hyperparameter set proposed in RawHash2. Still, since the original Sigmoni hyperparameter set achieved better results, we report Scrappie results in Sigmoni using the default Sigmoni hyperparameter set. We provide the hyperparameter setups for Scrappie in Sigmoni and RawHash2 frameworks in Table 1.

Table 1: Hyperparameter sets for Scrappie segmentation algorithm as defined in the Sigmoni and RawHash2 frameworks.

| **Hyperparameter** | **Sigmoni R9.4.1 & R10.4.1** | **RawHash2 R9.4.1** | **RawHash2 R10.4.1** |
|---|---|---|---|
| Window Length 1 | 3 | 3 | 3 |
| Window Length 2 | 6 | 9 | 6 |
| Threshold 1 | 4.30265 | 4.0 | 6.5 |
| Threshold 2 | 2.57058 | 3.5 | 4.0 |
| Peak Height | 1.0 | 0.4 | 0.2 |

#### S4 Scrappie RawHash2 and Sigmoni time comparison

Scrappie implementations used in the Sigmoni and RawHash2 frameworks are customized to improve downstream performance. These customizations include adjustments to the hyperparameters and thresholds of the Scrappie algorithm but do not modify the core algorithm itself. RawHash2, implemented in C++, integrates a C++ version of the Scrappie algorithm that processes signal chunks by first identifying positions corresponding to event borders and then computing normalized means of the signal events. In contrast, Sigmoni is primarily implemented in Python and uses a Python binding provided by the Uncalled4 framework to invoke the C++ Scrappie segmentation from Python. The segmentation function in Uncalled4 performs both event border detection and normalized event mean calculation in a single call.

Since the Scrappie implementation in RawHash2 and the one invoked by Sigmoni are functionally equivalent, we compare the times taken for border detection and event mean calculation in RawHash2 with the time taken to perform both steps within the Sigmoni framework. This comparison allows us to analyze the computational overhead introduced by bridging Python and C++ in Sigmoni, estimate the time Sigmoni spends solely on event border detection, and compare that with the time taken by Campolina to produce the same output.

Table 2: Time requirements for segmentation steps by RawHash2 and Sigmoni frameworks

| **Command** | **Mean execution time** |
|---|---|
| RawHash2 border detection | 14.84 |
| RawHash2 mean calculation | 11.89 |
| Sigmoni border + mean | 39.84 |

Table 2 shows the average time of five runs of segmentation steps in RawHash2 and Sigmoni frameworks. With the assumption that the main difference between the two implementations is the conversion between Python and C++ in Sigmoni, we see that the Sigmoni approach takes 13.10 seconds longer compared with the RawHash2 implementation. Furthermore, if we approximate the time Sigmoni would require for obtaining the border positions based on the execution time of border detection in RawHash2, and the previously mentioned time overhead from Python binding in Sigmoni, we see that obtaining border positions with Campolina, which is already implemented in Python, could be significantly more efficient compared with current Sigmoni segmentation framework.

#### S5 Supplementary figures

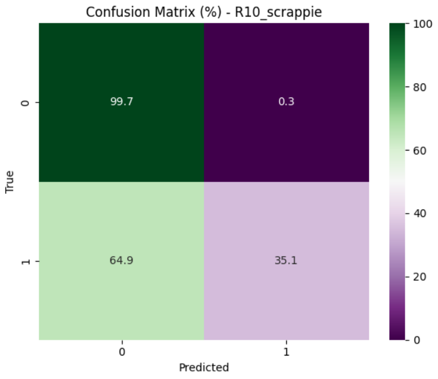


(a) Zymo binary Scrappie R10

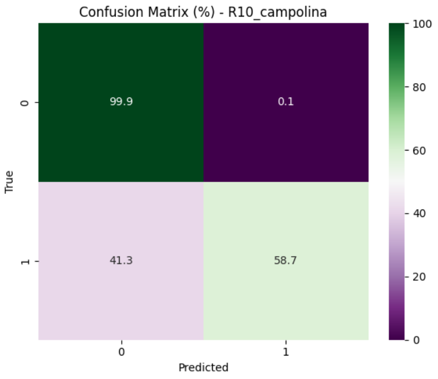


(b) Zymo binary Campolina R10

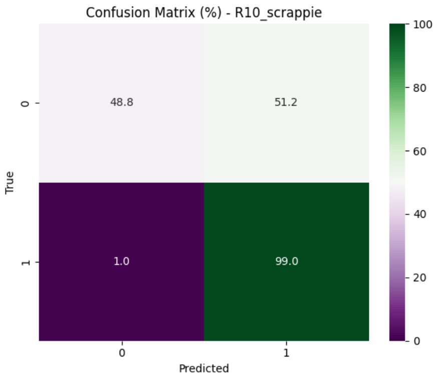


(c) Host depletion Scrappie R10

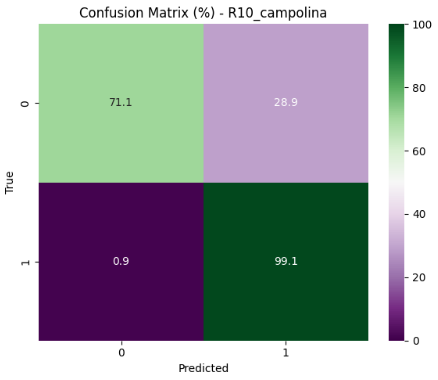


(d) Host depletion Campolina R10

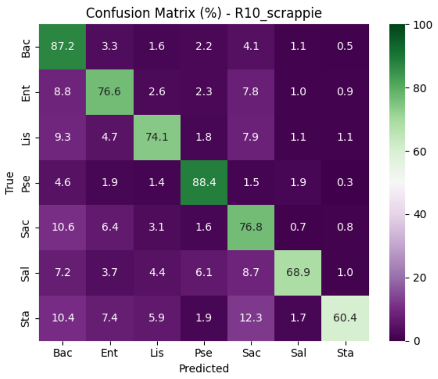


(e) Zymo multi Scrappie R10

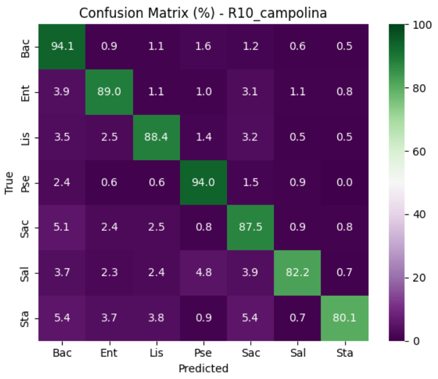


(f) Zymo multi Campolina R10

Figure 2: **Confusion matrices (percentage format) for the classification tasks on R10.4.1 datasets in Sigmoni framework.** **a)** and **b)** show results for the Zymo binary classification task using Scrappie, and Campolina segmentation strategy, respectively. Bacteria is labeled as class 0, Yeast is labeled as class 1. **c)** and **d)** show results for the Host depletion classification task using Scrappie, and Campolina segmentation strategy, respectively. Zymo is labeled as class 0, Human is labeled as class 1. **e)** and **f)** show results for the Zymo multiclass classification task, where *B. subtilis* is denoted *Bac*, *E. faecalis* is denoted *Ent*, *L. monocytogenes* is denoted *Lis*, *P. aeruginosa* is denoted *Pse*, *S. cerevisiae* is denoted *Sac*, *S. enterica* is denoted *Sal*, *S. aureus* is denoted *Sta*.

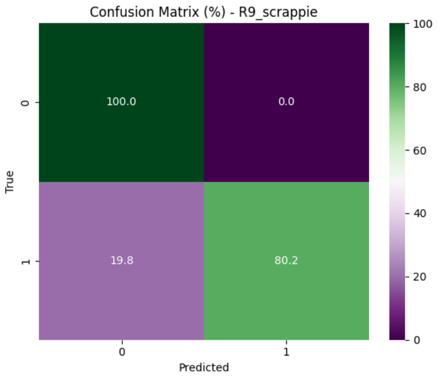


(a) Zymo binary Scrappie R9

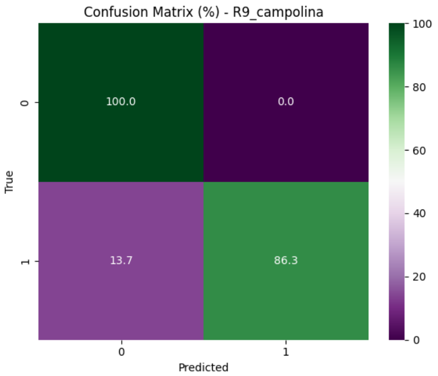


(b) Zymo binary Campolina R9

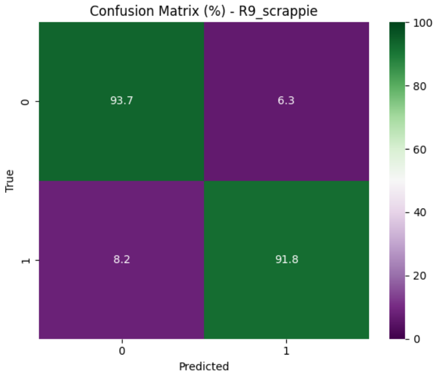


(c) Host depletion Scrappie R9

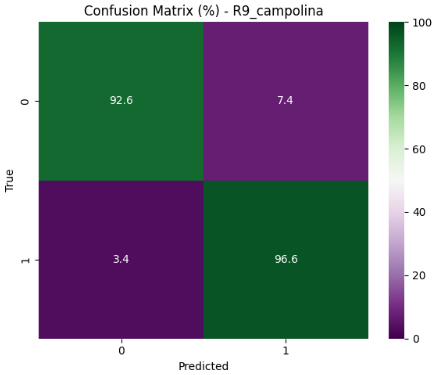


(d) Host depletion Campolina R9

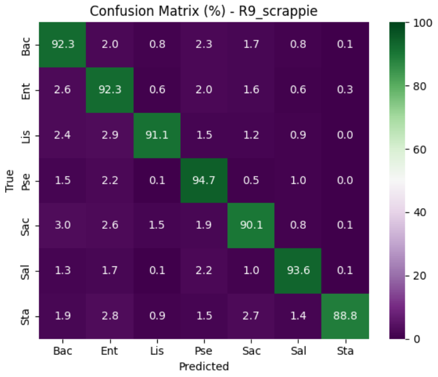


(e) Zymo multi Scrappie R9

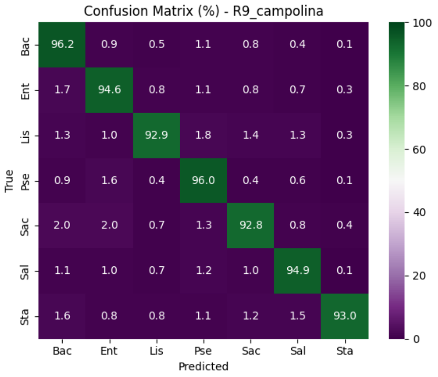


(f) Zymo multi Campolina R9

Figure 3: **Confusion matrices (percentage format) for the classification tasks on R9.4.1 datasets in Sigmoni framework.** **a)** and **b)** show results for the Zymo binary classification task using Scrappie, and Campolina segmentation strategy, respectively. Bacteria is labeled as class 0, Yeast is labeled as class 1. **c)** and **d)** show results for the Host depletion classification task using Scrappie, and Campolina segmentation strategy, respectively. Zymo is labeled as class 0, Human is labeled as class 1. **e)** and **f)** show results for the Zymo multiclass classification task, where *B. subtilis* is denoted *Bac*, *E. faecalis* is denoted *Ent*, *L. monocytogenes* is denoted *Lis*, *P. aeruginosa* is denoted *Pse*, *S. cerevisiae* is denoted *Sac*, *S. enterica* is denoted *Sal*, *S. aureus* is denoted *Sta*.

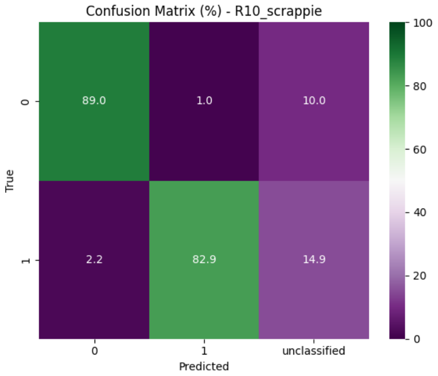


(a) Zymo binary Scrappie R10

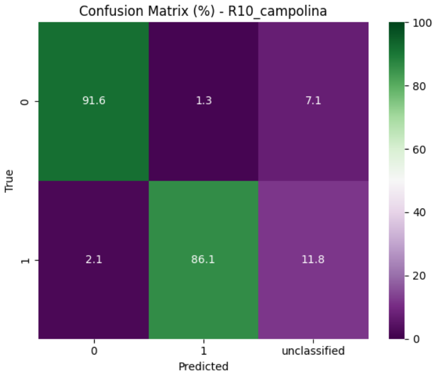


(b) Zymo binary Campolina R10

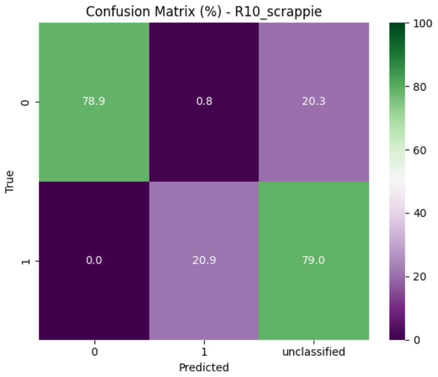


(c) Host depletion Scrappie R10

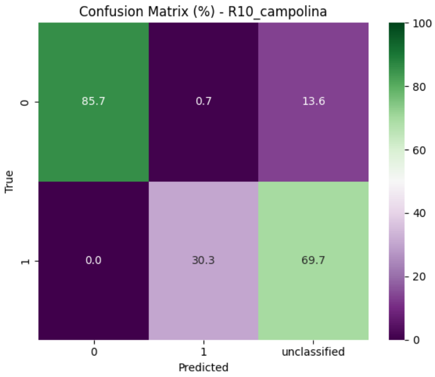


(d) Host depletion Campolina R10

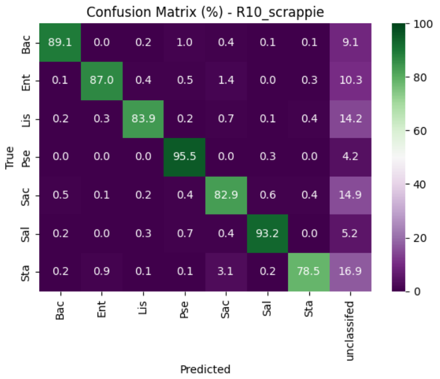


(e) Zymo multi Scrappie R10

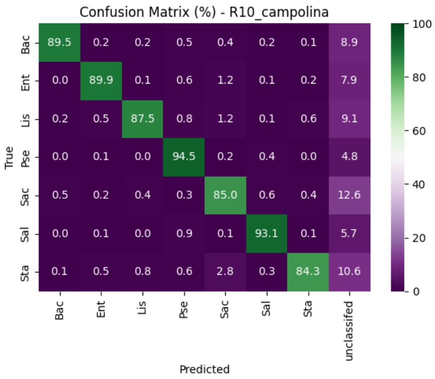


(f) Zymo multi Campolina R10

Figure 4: **Confusion matrices (percentage format) for the classification tasks on R10.4.1 datasets in RawHash2 framework.** **a)** and **b)** show results for the Zymo binary classification task using Scrappie, and Campolina segmentation strategy, respectively. Bacteria is labeled as class 0, Yeast is labeled as class 1. **c)** and **d)** show results for the Host depletion classification task using Scrappie, and Campolina segmentation strategy, respectively. Zymo is labeled as class 0, Human is labeled as class 1. **e)** and **f)** show results for the Zymo multiclass classification task, where *B. subtilis* is denoted *Bac*, *E. faecalis* is denoted *Ent*, *L. monocytogenes* is denoted *Lis*, *P. aeruginosa* is denoted *Pse*, *S. cerevisiae* is denoted *Sac*, *S. enterica* is denoted *Sal*, *S. aureus* is denoted *Sta*.

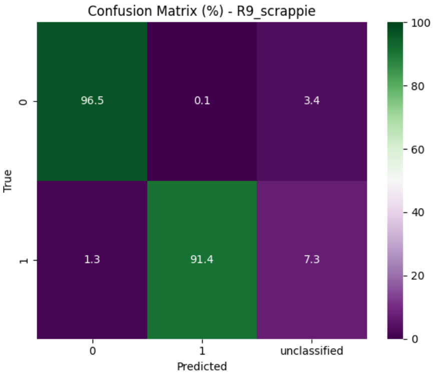


(a) Zymo binary Scrappie R9

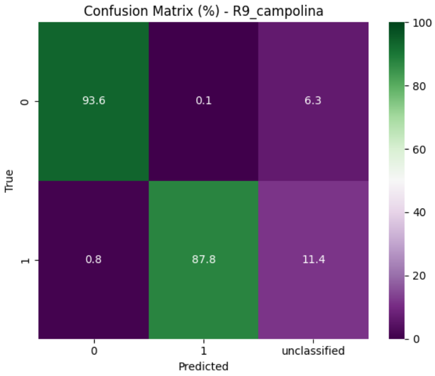


(b) Zymo binary Campolina R9

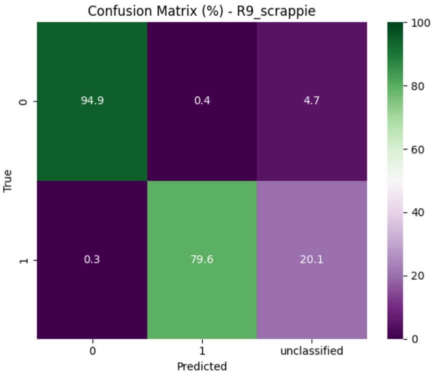


(c) Host depletion Scrappie R9

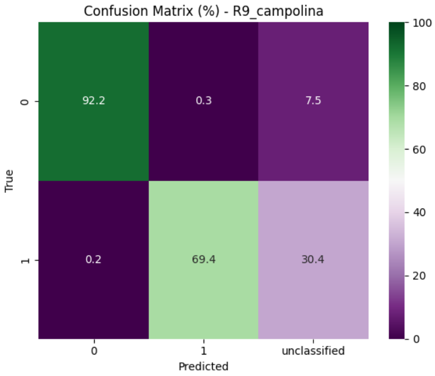


(d) Host depletion Campolina R9

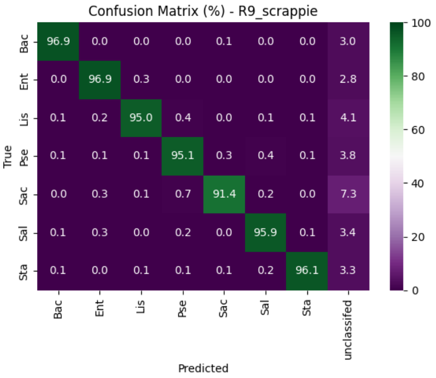


(e) Zymo multi Scrappie R9

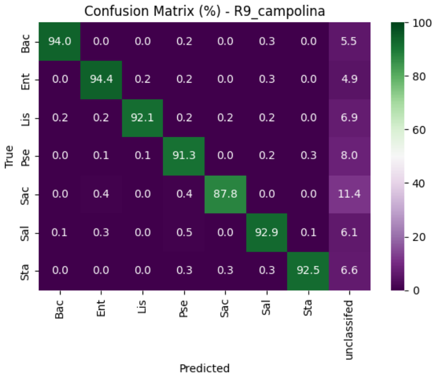


(f) Zymo multi Campolina R9

Figure 5: **Confusion matrices (percentage format) for the classification tasks on R9.4.1 datasets in RawHash2 framework.** **a)** and **b)** show results for the Zymo binary classification task using Scrappie, and Campolina segmentation strategy, respectively. Bacteria is labeled as class 0, Yeast is labeled as class 1. **c)** and **d)** show results for the Host depletion classification task using Scrappie, and Campolina segmentation strategy, respectively. Zymo is labeled as class 0, Human is labeled as class 1. **e)** and **f)** show results for the Zymo multiclass classification task, where *B. subtilis* is denoted *Bac*, *E. faecalis* is denoted *Ent*, *L. monocytogenes* is denoted *Lis*, *P. aeruginosa* is denoted *Pse*, *S. cerevisiae* is denoted *Sac*, *S. enterica* is denoted *Sal*, *S. aureus* is denoted *Sta*.
